# Phase relations of interneuronal activity relative to theta rhythm

**DOI:** 10.1101/2023.05.11.540330

**Authors:** Ivan Mysin

## Abstract

The theta rhythm plays a crucial role in synchronizing neural activity during attention and memory processes. However, the mechanisms behind the formation of neural activity during theta rhythm generation remain unknown. To address this, we propose a mathematical model that explains the distribution of interneurons in the CA1 field during the theta rhythm phase. Our model consists of a network of seven types of interneurons in the CA1 field that receive inputs from the CA3 field, entorhinal cortex, and local pyramidal neurons in the CA1 field. By adjusting the parameters of the connections in the model. We demonstrate that it is possible to replicate the experimentally observed phase relations between interneurons and the theta rhythm. Our model predicts that populations of interneurons receive unimodal excitation and inhibition with coinciding peaks, and that excitation dominates to determine the firing dynamics of interneurons.

## 1 Introduction

The hippocampus is a brain structure that plays a key role in the processes of attention and memory. To process information, neural ensembles in the hippocampus need to be synchronized with rhythms. The main rhythm that organizes the neural activity of the hippocampus during cognitive tasks is the theta rhythm (4-12 Hz) [10, 12, 63]. Almost all hippocampal neurons are modulated by theta rhythm [45]. This is expressed in the fact that each population has a theta rhythm phase, in which the probability of discharges of its neurons is maximal. Modulation of neuronal activity by rhythm makes it possible to synchronize different areas of the hippocampal formation during information processing [24, 48, 50]. Also, the theta rhythm provides an ordered structure of the place cell activity, due to phase precession [9, 31]. Thus, understanding the mechanisms of theta rhythm formation is the most important problem of neuroscience.

Several theoretical studies have investigated the formation of phase relations between neurons of the CA1 field and theta rhythm [6, 47]. Despite the detailed nature of these models, the authors were unable to reproduce the form of distribution of most types of interneurons in the theta rhythm phase. The main problem in constructing a model of the distribution of neurons by theta rhythm phases is to identify the mechanism of stabilization of antagonistic relationships between different populations of interneurons. For example, parvalbumin-containing (PV) and cholecystokinin-containing (CCK) basket neurons inhibit each other and fire at opposite phases of the theta rhythm [7, 40, 52, 55, 60]. It is difficult to choose the parameters of connections when these neurons form a stable antiphase activity. Most often, one of the populations completely inhibit the other. When considering a larger number of interneuron populations, the problem becomes much more complicated [6, 47]. We hypothesized that short-term synaptic plasticity stabilizes the structure of interneuronal activity. In a recent study, the authors of the project Hippocampome.org provide a large meta-analysis of data on short-term synaptic plasticity in the hippocampus [46]. We used the estimates of this work as the basis of our model. We have considered the mechanism of formation of phase relations for populations of interneurons of the CA1 field. The CA1 field was chosen because there is the greatest amount of data on interneuron activity for this region of the hippocampus. We modeled 7 populations of interneurons: PV and CCK basket, axo-axonal, OLM (oriens-lacunosum moleculare), Ivy, neurogliaform and bistratified neurons. These populations were selected because the phase relationships for them are described [25, 39, 55, 56, 60]. Neurons in these populations make up about 70 % of all interneurons in the CA1 field [7]. We also took into account inputs from pyramidal neurons of the CA3 and CA1 fields, as well as from neurons of the 3rd layer of the entorhinal cortex. We were able to select the parameters of neurons and connections in the model and found a good description of the mechanisms of phase relations.

## 2 Materials and methods

### 2.1 Models of neurons

Monte Carlo simulations were provided with equations:

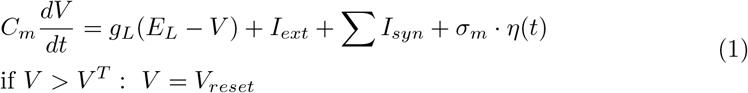

here *C*_*m*_ is the membrane capacitance, *V* is the membrane potential, *g*_*L*_ is the leak conductance, *E*_*L*_ is the leak reverse potential, *I*_*ext*_ is the external current and *I*_*syn*_ is the synaptic current. *σ*_*m*_ is the standard deviation of noise *η ∼ N* (0, 1). For all neurons in all simulations we used follow values: *C*_*m*_ = 1 *μF/cm*^2^, *g*_*L*_ = 0.1 *mS/cm*^2^, *E*_*L*_ = *−*60 *mV, V* ^*T*^ = *−*50 *mV, V*_*reset*_ = *−*90 *mV*, 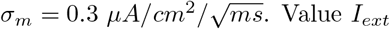. Value *I*_*ext*_ is optimized for each population. After generating the action potential, the refractory = 3 *ms*

Population frequency for Monte Carlo simulations:

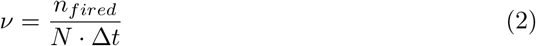

where *n*_*fired*_ is the number fired cell at each time step, *N* = 4000 is the full number of neurons in population, Δ*t* = 0.1 *ms* is the integration step. We used stochastic Heun method from brian2 package for Monte Carlo simulations [57].

### 2.2 Synapse model

Synapses were simulated with Tsodyks-Markram model [46, 58]:

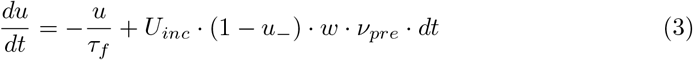

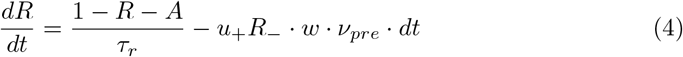

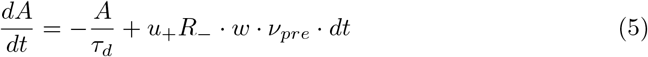

*U*_*inc*_ is the increment of *u* produced by a spike. *u*_*−*_, *x*_*−*_ are variables just before the arrival of the spike, and *u*_+_ refers to the moment just after the spike. From the first equation, *u*_+_ = *u*_*−*_ + *U*_*inc*_ .(1 *−u*_*−*_). Variables *A* and *R* means the activated deactivated, and recovered states respectively. After each presynaptic spike, an instantaneous shift occurs from recovered to activated state. The amount of shift is determined by *u*. The active resources then decay to the deactivated state by the decay time constant *τ*_*d*_. Since synaptic resources are limited, the more resources stay in the deactivated state, the more a synapse is depressed. Synaptic resources exponentially recover from depression with the recovery time constant *τ*_*r*_. *w* is normalization coefficient, it makes sense of the density of connections. *ν*_*pre*_ is firing rate of presynaptic population. In Monte Carlo simulations *ν*_*pre*_ is defined by equation 2, in CBRD simulations *ν*_*pre*_ is defined by equation 9. For each connection parameters *U*_*inc*_, *τ*_*d*_, *τ*_*f*_, *τ*_*f*_, *w, g*_*syn,max*_ are optimized. The synaptic current:

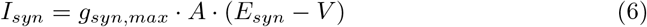

where *g*_*syn,max*_ is maximal synaptic conductance, *E*_*syn*_ is reversal potential for synaptic current. The connection probability in Monte Carlo simulations is 0.5.

### 2.3 CBRD approach

CBRD approach simulates the distribution (*ρ*) of neurons in space-times after spike the generation (*t*^*\**^) [14–16]. The function *ρ*(*t, t*^*\**^) is described by the transfer equation:

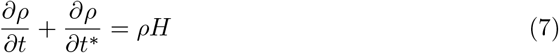

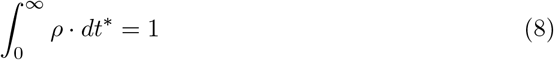

*H* is probability of spike generation (eq. 11). Population firing rate is defined by

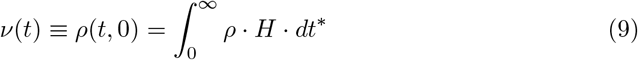

Neuron model modified:

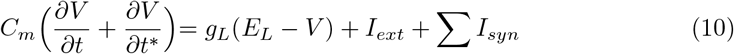

Equations of CBRD approach:

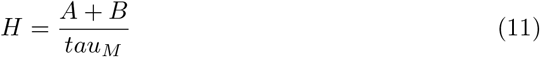

*tau*_*M*_ is time of membrane:

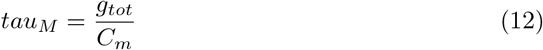

*C*_*m*_ is capacity of membrane, *g*_*tot*_ is total conductance of all channels, it is sum of *g*_*L*_ and all synaptic conductances.

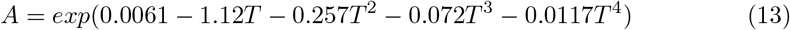

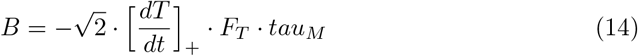

[]_+_ is Heaviside function.

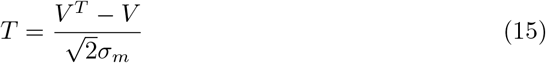

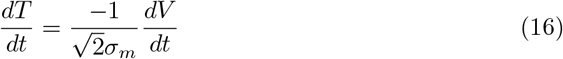

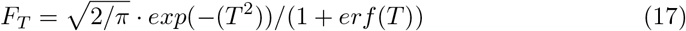

In these equations *V* is membrane potential, *σ*_*m*_ is the sigma of noise, *V* ^*T*^ is the threshold for spike generation, *erf* is Gauss error function. All values for model are same as for Monte Carlo simulations.

### 2.4 Numerical methods for CBRD simulations

In general, the equations can be written as

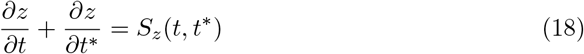

where *z*(*t, t*^*\**^) is one of the functions *ρ* or *V, S*_*z*_(*t, t*^*\**^) is the source term of the equation.

Numerical scheme were taken from [28]. Numerical scheme is

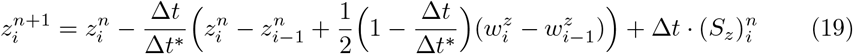

*n* is index in time, *i* is index in space.

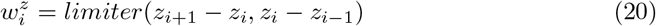

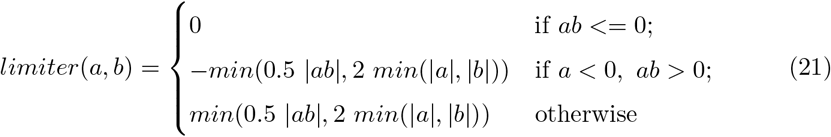

### 2.4.1 Bound conditions

Bound values 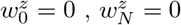. Firing rate (left bound for *ρ*) is calculated by

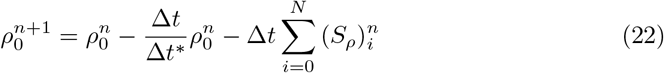

Left bound for *V* :

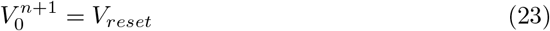

Right bound for *ρ*:

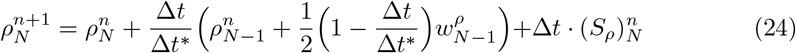

Right bound for *V* :

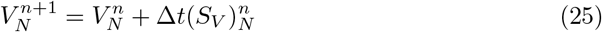

Δ*t* = 0.1 *ms*, Δ*t*^*\**^ = 0.5 *ms, N* = 400 is the number of spatial states. All simulations using the CBRD approach were performed with the TensorFlow package (version 2.10.0). All gradients were calculated using automatic differentiation [2].

### 2.5 Target firing rates, inputs and loss function

Firing rate of inputs and target firing rate for simulated populations:

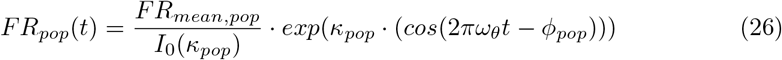

where *FR*_*pop*_(*t*) is population firing rate in time in spikes/second. *FR*_*mean,pop*_ is mean population firing rate (sp./sec). *kappa*_*pop*_ is a measure of concentration of von Mises distribution, *I*_0_(*κ*) is a zero order Bessel function. *ω*_*θ*_ = 7 *Hz* is frequency of theta rhythm, *ϕ*_*pop*_ is peak phase of population firing. Parameters for all populations are presented in table 1.

**Table 1.**
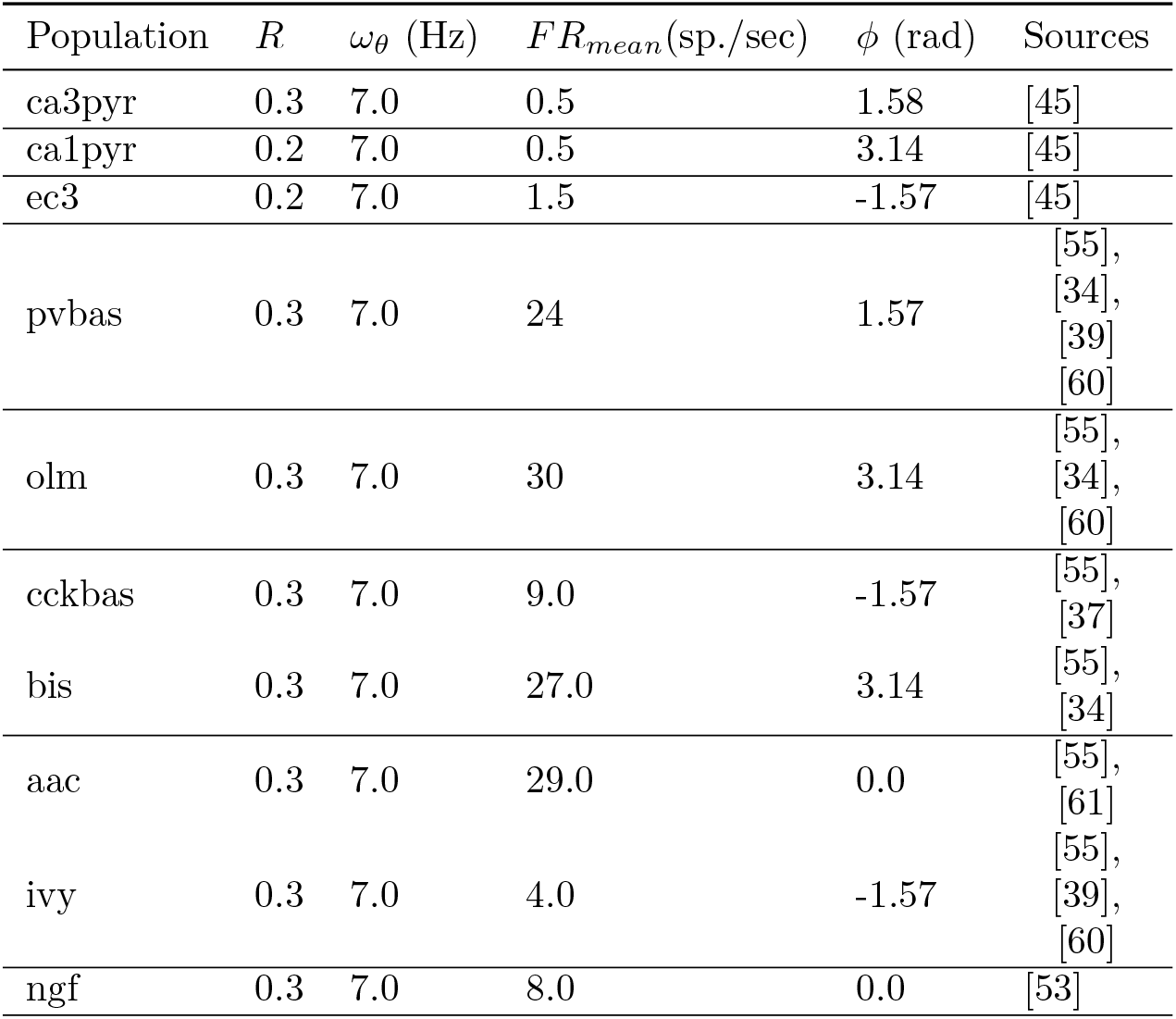
Parameters of the target function.

*κ* is calculated from *R* with follow approximation [1]):

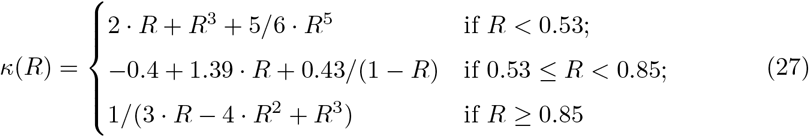

Artificial inputs in Monte Carlo simulations are modeled as Poisson generators with rate given by the equation 26.

The loss function for estimating the discrepancy between the simulation and the target function:

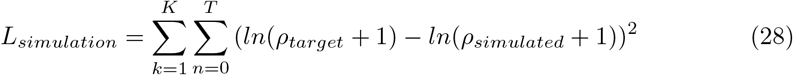

Summation is carried out by time and by all populations of neurons.

The optimized parameters must be within certain bounds. In particular, all parameters of synapses should be positive, and *U*_*inc*_ should not exceed one. We have added barrier terms to the loss function

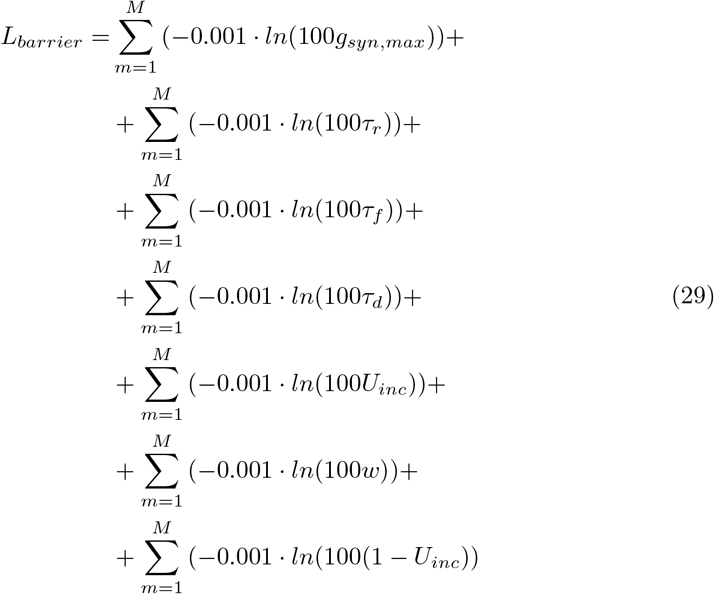

Summarization is carried out for all synapses in the model.

Full loss function for optimization:

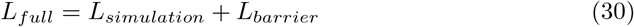

We have used Adam optimizer with standard parameters: learning rate = 0.001,

*β*_1_ = 0.9, *β*_2_ = 0.999 [35].

## 3 Results

### 3.1 Optimization of the model

We considered a network of biologically plausible networks consisting of 7 populations of interneurons. The graph of connections is complex and contains 49 connections (fig. 1) [46]. Excitatory inputs were not modeled as neural populations, but were taken into account artificial generators (functions from time). Each synapse is described by 6 parameters (eq. 3, 4, 5). We fit all these parameters. Also, the *I*_*ext*_ parameter for each neuron population was fit in simulations (eq. 1). Thus, we needed to tune 301 model parameters. In our study, we applied optimization techniques to find the model parameters that best describe the experimental data. Our approach contains elements of novelty, so we will begin the description of the results with a brief description of the formulation and solution of the optimization problem.

**Figure 1.**
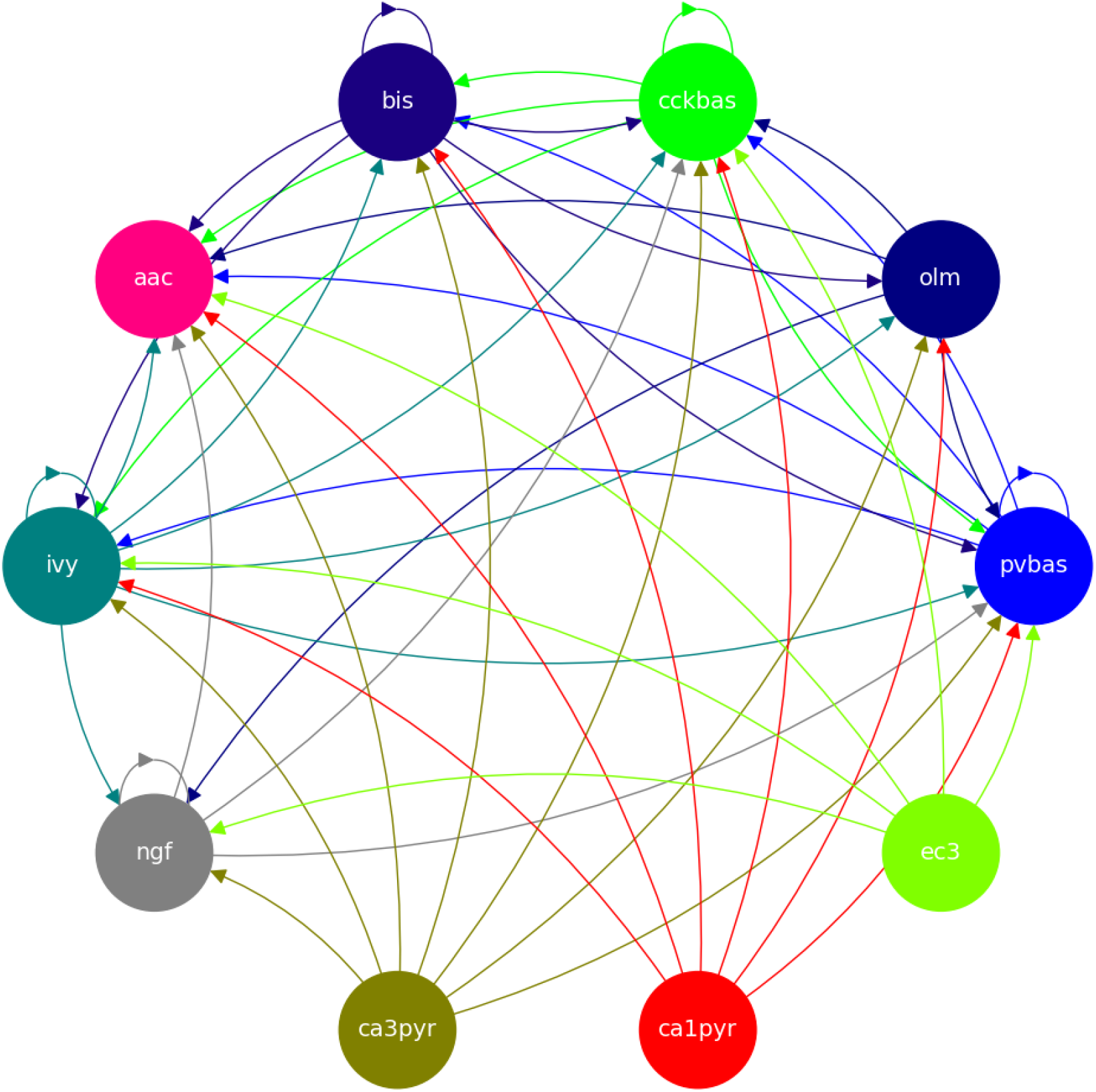
Graph of connections between neural populations. Note: ca3pyr - pyramidal neurons of the CA3 field, ca1pyr - pyramidal neurons of the CA1 field, ec3 - principal neurons of the 3 layer of the entorhinal cortex, pvbas - PV basket cells, olm - Oriens-lacunosum molecular cells, cckbas - CCK basket cells, Ivy - Ivy cells, ngf - neurogliaform cells, bis - bistratified cells, aac - axo-axonal cells.

We have introduced a target activity function for each population (eq.26). This function describes the population firing rate of each group of neurons over time. The target function takes into account the average firing rate and theta rhythm modulation. Theta rhythm modulation is determined by the mean phase (circular average of the phases of neuronal discharges) and the phase variation (ray length - R). The population firing rate of excitatory inputs were described by similarly functions (eq. 26).

We aimed to create a model that would describe the entire form of distribution of neuronal activity over the theta rhythm phase. We have used the population approach for modeling, since it allows us to describe the population firing rate. As a population approach, we chose the CBRD (conductance base refectory density) method [14–16]. Direct simulation of the firing rate makes it possible to estimate the gradient of the loss function from the model parameters. Estimating the gradient of the loss function makes it possible to apply gradient descent methods to optimize model parameters (eq. 30). All simulations using the CBRD approach were performed with the TensorFlow package, gradients were calculated using automatic differentiation [2].As initial conditions for optimization, we used the average values of the parameters described in [46].

This approach has advantages and disadvantages. On the one hand, population models are only applicable to simple models of neurons. In our work we used the LIF (leaky integrate-and-fire) model (eq. 1 On the other hand, the use of gradient descent allows optimization over a much larger number of parameters than gradient free methods.

### 3.2 Description of the model results

Fig. 2 shows the optimization results. For each population, the target function, the results of simulation using the CDRD approach, and the results of Monte Carlo simulation with optimal parameters are presented. The model under consideration can approximate the target functions for all populations of interneurons with significant accuracy. The model describes all the required characteristics of the experimental data: the average spike rate, the average phase of the theta rhythm, and the ray length (R). We also note a good agreement between the results of Monte Carlo simulations and simulations using the CBRD approach. In the next step, we looked at how the model works. Fig. 3 shows the dynamics of synaptic conductivities in each neuron population. Both excitatory and inhibitory conductivities have an unimodal distribution over the theta cycle for all neuron populations, except neurogliaform cells. An unexpected result is that the peaks of the excitatory and inhibitory inputs coincide. This effect is observed for all populations of neurons. The ratio of the total excitatory to inhibitory conductivity remains approximately constant over the theta cycle and ranges from 0.7 to 2.4 for different populations. Since the reversion potential for AMPA receptors (0 mV) is much further from the resting potential than the reversion potential of GABA-A receptors (−75 mV). Excitatory currents dominate over inhibitory ones and determine the dynamics of neuronal discharges. For all neuron populations except axo-axonal ones, the peak of excitatory input is located near the peak of firing rate. The dynamics of axo-axonal neurons in the model is determined by the external current (Table **??**). All optimal parameters are given in Supplementary materials.

**Figure 2.**
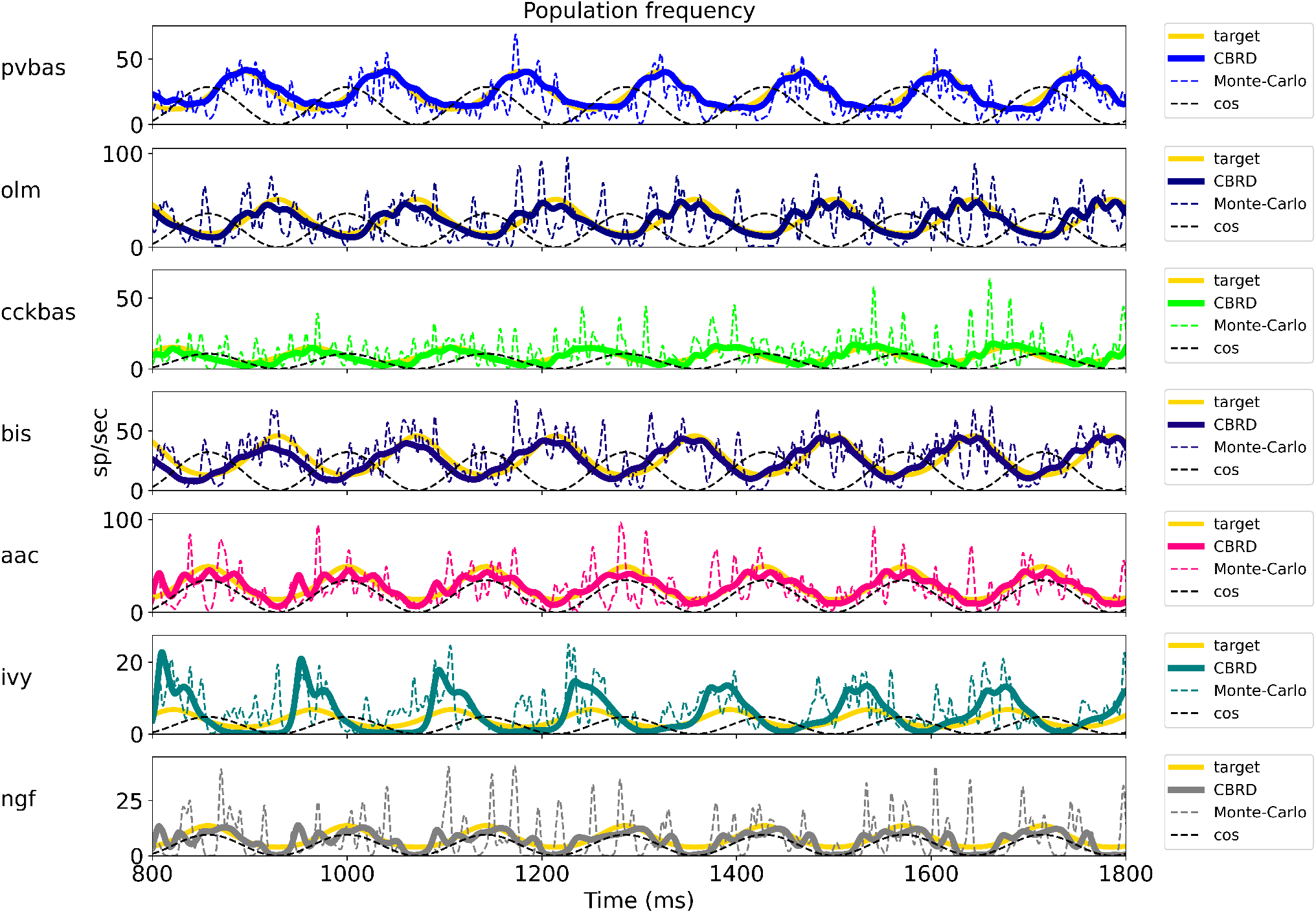
Results of model optimization. For each population, plots show the target function, the population spike rate obtained with the CBRD approach and using Monte Carlo simulation. One second of simulation is shown after stabilization of the model dynamic mode. The notation of neurons is similar to fig. 1

**Figure 3.**
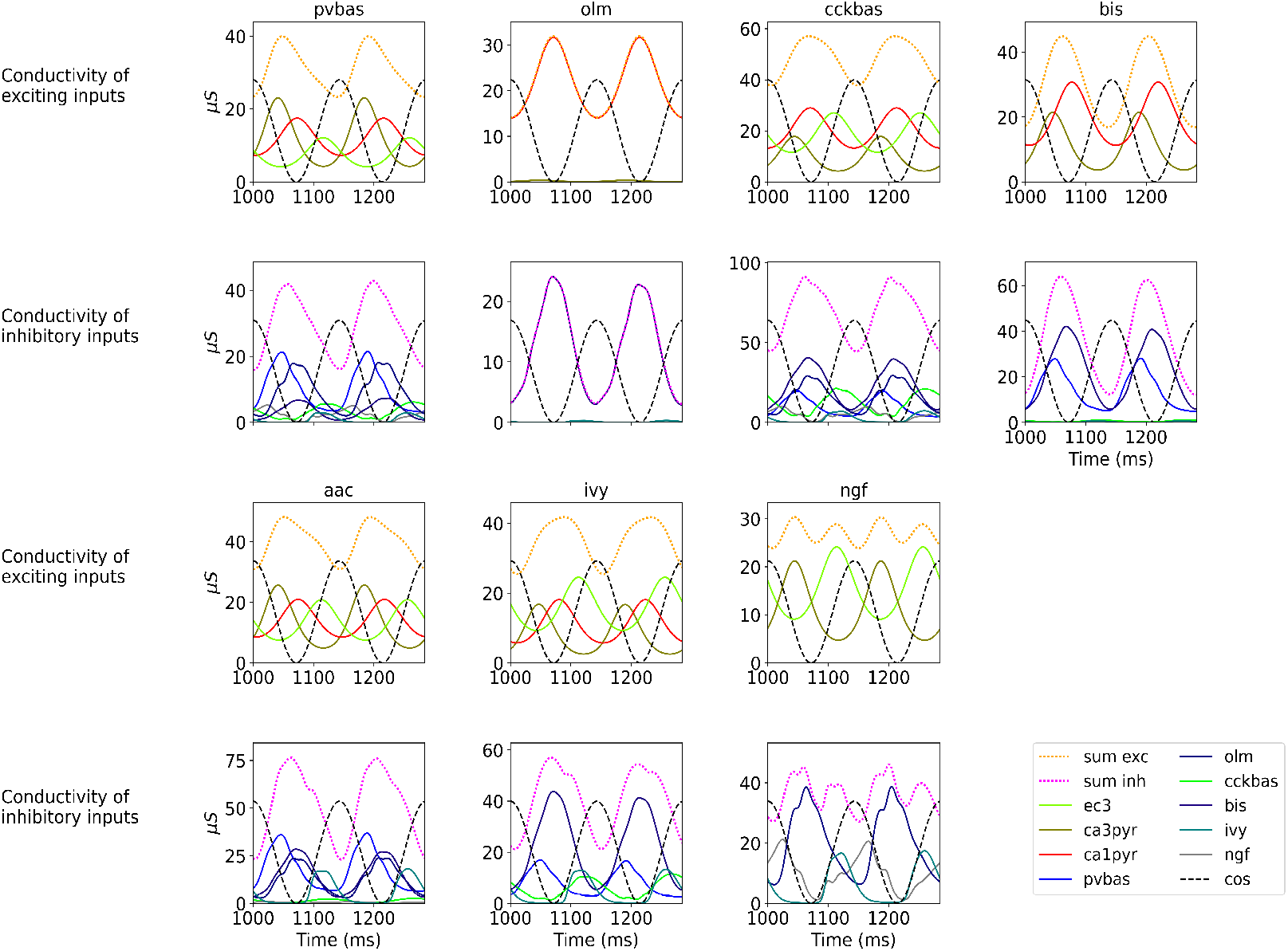
Synaptic conductivities. Two theta cycles of the simulation using the CBRD approach are shown. Excitatory (upper series of plots) and inhibitory (lower series of plots) conductivities are shown for each population. The color indicates presynaptic populations. Each plot shows the sum of conductivities, “sum exc” and “sum inh” notes sum of excitatory and inhibitory conductivities, respectively. The notation of neurons is similar to fig. 1

We performed optimization for a similar network with synapses without short-term plasticity. The results are presented in the Supplementary materials. For a part of neural populations (pvbas, olm, bis, aac), it fits the parameters to describe phase relations. However, the remaining populations are completely silent (ivy, ngf, cckbas). These results show that short-term plasticity is a necessary element of the model for reproducing phase relations.

### 3.3 Generalizing power of the model

At the last stage of research, we tested the stability of the found solution at other theta rhythm frequencies. We searched for optimal parameters at a theta rhythm frequency of 7 Hz. Fig. 4 shows the results of simulations with varying theta rhythm frequency. Fig. 5 shows a raster plots for a Monte Carlo simulation with a theta rhythm frequency of 5 Hz. In this series of computational experiments, we changed the frequencies of the input populations. The model parameters were used that were found earlier. Over the entire frequency range (4-12 Hz), the shape of the distribution of neuronal activity over the theta rhythm phase is preserved. The theta rhythm frequency is a parameter that varies greatly in animal experiments, for example, it correlates with the animal’s running speed [29, 33, 41, 43]. The conservation of the phase relations of interneuron activity with a change in the theta rhythm frequency is an important property of our model. These experiments can be considered as analogous to cross-validation used in machine learning. In the language of machine learning, we can say that our model shows good quality on the “test set”, i.e. on samples that did not participate in the “training”.

**Figure 4.**
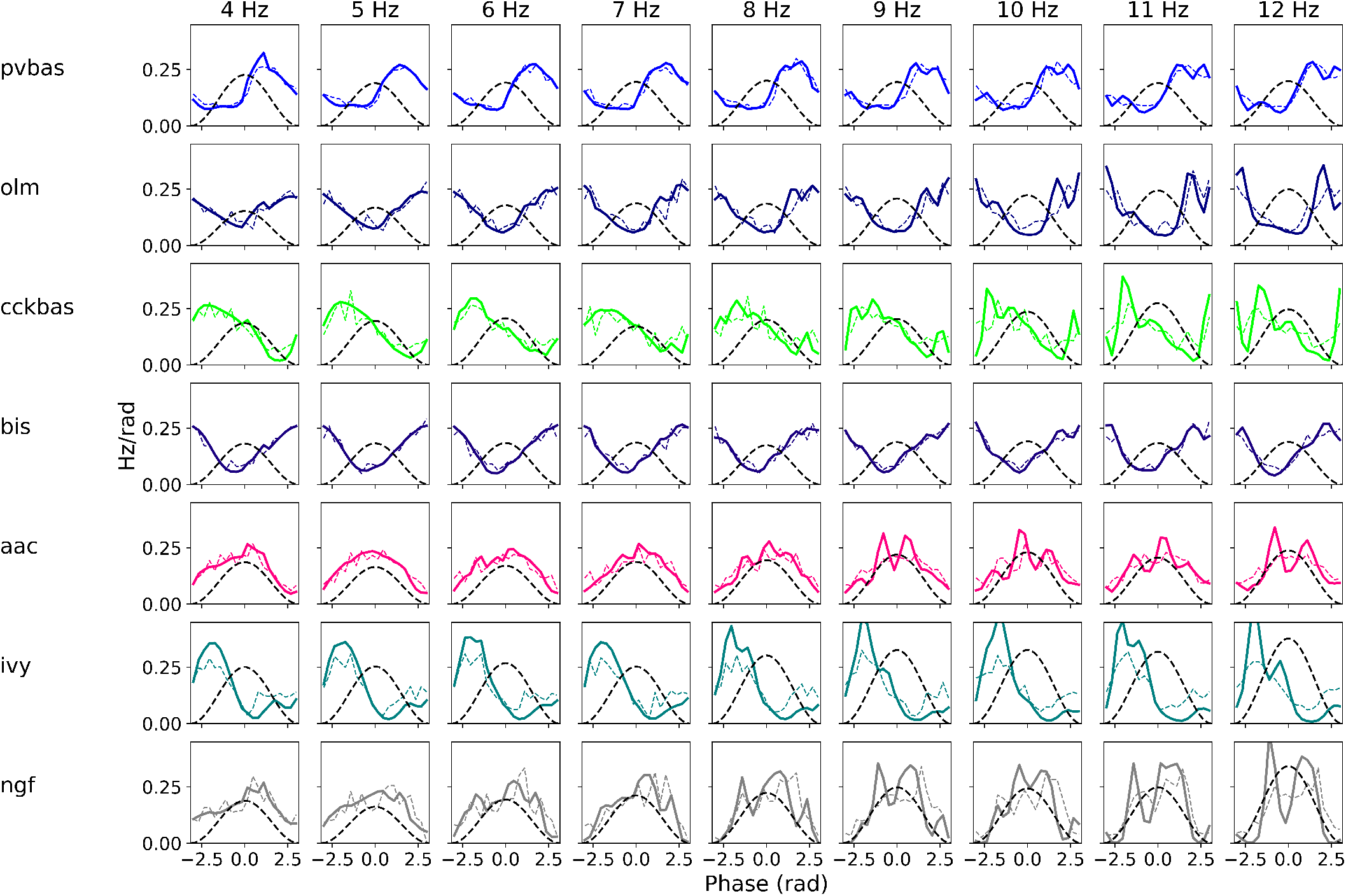
Phase relations. Each plot shows the distribution of activity of neuronal populations by theta rhythm phase. The frequency of the theta rhythm was set in the equations of the exciting inputs. All simulations were carried out with optimal parameters found at a theta rhythm frequency of 7 Hz. Simulations were carried out using the CBRD approach (thick lines) and Monte Carlo (dashed line). The notation of neurons is similar to fig. 1

**Figure 5.**
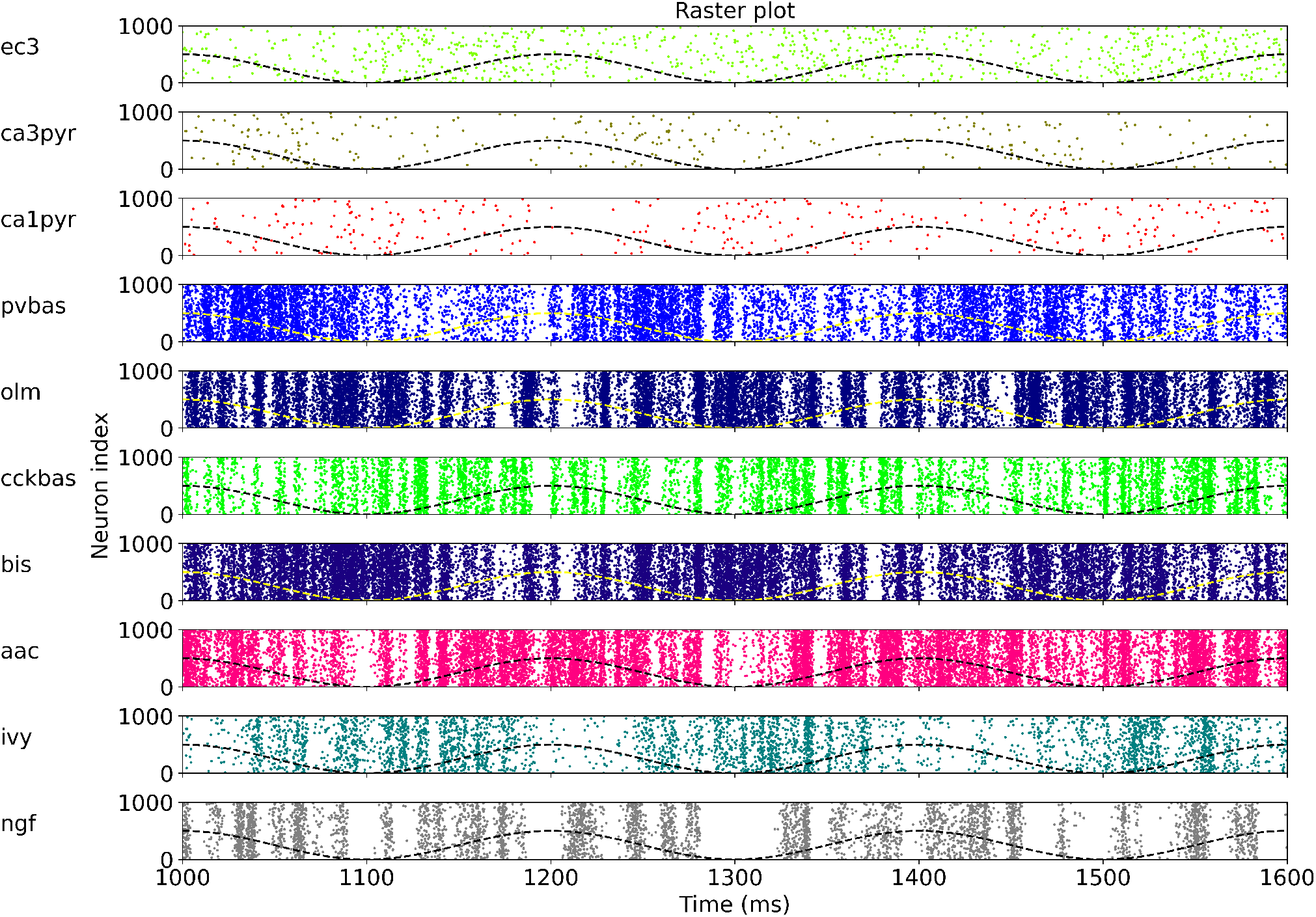
Raster plots for Monte Carlo simulation with the theta rhythm frequency of 5 Hz. For each population only 1000 neurons are shown, but it was simulated 4000. Dashed line is reference cos. The notation of neurons is similar to fig. 1

## 4 Discussion

### 4.1 Assumptions and limitations of the model

Our results are based on several key assumptions. We used the von Mises function to describe the inputs and target activities for interneurons (eq. 26). These assumptions of the model are based on experimental data that more than 85 % of pyramidal neurons in the CA3 and CA1, 3 fields of the EC layer are modulated by theta rhythm [45]. The proportion of interneurons of the CA1 field modulated by the theta rhythm is 95 % [45]. The degree of phase modulation R for all hippocampal neurons is in the range of 0.2-0.4; we used R = 0.3 as the mean estimate. Binding phases for exciting inputs and local interneurons of the CA1 field (table 1) have been measured in several experimental studies and are reliable [36, 45, 55, 56]. Our model takes into account the average discharge frequencies of input populations and interneurons (table 1). We selected the values for simulations as the mean and median values measured in the literature. The exact values of the mean firing rate are not important in our context, since the discharge frequency of the presynaptic population is linearly related to the weight of the synaptic connection with the postsynaptic population. In other words, a decrease of the firing rate of the presynaptic population can be compensated by an increase in synaptic conduction in postsynaptic neurons. We used a homogeneous representation of each population of interneurons. Recent studies show that interneurons are involved in the formation of neural ensembles, i.e. their activity is modulated by external stimuli [26]. However, a study of simultaneous registration of numerous interneurons of different classes shows that cells of all classes are active simultaneously for times of hundreds of milliseconds [27]. Thus, it is reasonable to assume that a significant proportion of neurons in each population are active at the same time. Note that the effect of simultaneous activity of interneurons of different populations is observed, despite the inhibitory connections of populations between each other [7]. Our model reproduces this effect well. Due to the homogeneity of each population, we used the same number of neurons in direct simulations. Interneurons are unevenly distributed across classes. For example, population of Ivy neurons are in 6 times larger than axo-axonal cells [7]. A representative representation of neurons would not improve the accuracy of the model, but it would increase computational costs. We did not take into account the input to the model from the medial septum. Recent studies show that the strongest rhythmic input from the septum goes to the interneurons of the CA3 field and the entorhinal cortex [32, 59, 62]. Other data demonstrate the importance of input from the CA3 field and the entorhinal cortex to the CA1 field for theta rhythm in it [44, 67]. Together, these results lead to the hypothesis that the theta rhythm in the CA1 field results from secondary rhythmic inputs, rather than direct input from the medial septum. There are two significant limitations of our model. The first is usage the LIF model of neurons. Most types of interneurons have slow potassium channels [6, 38] which can make a significant contribution to the frequency of neuronal discharges due to spike-frequency adaptation. OLM neurons have pronounced H-currents [54] which can also make a significant contribution to the impulsing of these neurons due to depolarization or resonant properties [3]. The usage of the LIF model is dictated by a limitation of the population approach. We did not explicitly simulate pyramidal neurons of the CA1 field due to the limitations of the population method. Different classes of interneurons have entrances to different compartments of pyramidal cells. The correct inclusion of pyramid neurons in the network requires the use of multicompartment models to describe them. The problem of finding optimal synaptic inputs to pyramidal neurons can be solved separately.

The second limitation is that we did not take into account all populations of interneurons. This limitation is based on the weak knowledge of other populations of interneurons. The lack of data makes it impossible to include other populations in the model. Accounting for more populations can change the balance of inhibition and shift its peak.

### 4.2 Comparison with other models

The literature presents several attempts to explain the phase relations between neurons and theta rhythm. The key idea of the research is to determine the optimal structure of the input from the medial septum and excitatory inputs [6, 19–21, 47, 49]. Our results do not negate the contribution of external inputs, but rather dismantle an additional mechanism for stabilizing phase relations. Models with a few interneuron populations give a good approximation of the distribution of neurons in theta rhythm phases due to input from the medial septum [19, 49]. However, with an increase in the number of interneuron classes in models, the quality of the approximation of phase relations decreases [6, 21, 47]. Although in the cited papers, the authors used a different input structure. We believe that taking into account the detailed structure of the inputs to the CA1 field in the model is not sufficient to explain the phase relations of interneurons. In this paper, we have shown that the short-term plasticity of synapses between populations of interneurons may be the missing component in the stabilization of phase relationships.

### 4.3 Comparison of simulation results with experimental data

The main result of our work is an unimodal distribution of excitation and inhibition with coinciding peaks for most types of interneurons. Checking this fact is a complex experimental task. In the literature, there is a study of synaptic currents on PV basket and OLM neurons during theta rhythm generation in hippocampal slices [30]. The authors also found that unimodal excitation and inhibition coincided in time. The peak of inhibition occurred 12 ms after the peak of excitation for PV basket neurons, and 7 ms after the peak of arousal for OLM neurons [30]. We used experimental data in vivo as the basis for our model therefore, comparison with data on hippocampal preparations is limited.

### 4.4 Prospects for applying population model optimization

Several attempts have been made in the literature to optimize networks of spiking neurons [42, 51, 65, 66]. However, these works are aimed at finding practical applications in the field of data analysis, not brain modeling. These approaches are not directly applicable for building a model in neuroscience. The activity of real individual neurons is noisy. The vast majority of experimental phenomena are obtained as a result of averaging the activity of one or several neurons, depending on external stimuli or internal state. The phase relations of neuronal activity are one of many examples. Another example is place cells - the distribution of neuronal activity depending on the position of the animal, phase precession is the distribution of neuronal activity depending on the position of the animal and the phase of the theta rhythm, etc. Population models describe physiological phenomena in the language of distributions. Modeling complex systems requires numerous equations and parameters. Projects such as the Human Brain Project or the Hippocampus Project aim to model the brain by collecting and organizing parameters for equations [8, 64]. Optimization of population models can well complement the processes of collecting parameters. The development of the mathematical apparatus of population models may be one of the reasons for a breakthrough in the construction of large-scale models of the brain.

### 4.5 General remarks

Our model shows a possible mechanism for the formation of phase relations between interneurons and theta rhythm in the CA1 field. We assume that a similar mechanism operates in other areas of the hippocampal formation. The establishment of phase relationships among interneurons leads to the coupling of principal neurons to the phase of the theta rhythm. Rhythmic inputs to the principal neurons creates oscillations of local field potential [11, 22]. This, in turn, synchronizes the regions of the hippocampus with each other through the projections of principal neurons, which is necessary for the transmission of information [4, 48]. Synchronization of hippocampal regions creates an capability for the formation of phase precession [9, 23, 67]. Another result of the synchronization of different areas of the hippocampus is the emergence of coupling of theta and gamma rhythms. For example, a slow gamma rhythm is formed in a feedback loop between PV basket and principal neurons [13, 18]. In the CA1 field, excitation from the CA3 field is necessary for the formation of a slow gamma rhythm [17, 23]. PV basket neurons CA1 have a peak of discharges in the descending phase of theta rhythm, the input from field CA3 falls into the same phase [5, 45]. Therefore, a model that reproduces phase relationships with respect to the theta rhythm will have a good predictive ability with respect to other phenomena associated with the theta rhythm. Identification of the mechanisms of formation of the dynamics of the activity of interneurons during the generation of the theta rhythm is important for understanding the processing of information and the formation of hippocampal rhythms.

