## Supplementary Materials for "Phase relations of interneuronal activity relative to theta rhythm"

### Supplementary Materials (”Phase relations of interneuronal activity relative to theta rhythm)

#### 1 Optimization results of the model with non-plastic synapses

Nonplastic synapses simulated with equation:

$$\tau_1 \cdot \tau_2 \cdot \frac{d^2 g_s}{dt^2} + (\tau_1 + \tau_2)\frac{dg_s}{dt} = g_s \cdot w \cdot \nu_{pre} \tag{1}$$

The synaptic current:

$$I_{syn} = g_{syn,max} \cdot g_s \cdot (E_{syn} - V) \tag{2}$$

$g_{syn,max}, \tau_1, \tau_2, w$ were optimized. The barrier term in the loss function has been modified:

$$\begin{aligned} L_{barrier} = &\sum_{m=1}^{M} (-0.001 \cdot ln(100 g_{syn,max})) + \\ &+ \sum_{m=1}^{M} (-0.001 \cdot ln(100\tau_1)) + \\ &+ \sum_{m=1}^{M} (-0.001 \cdot ln(100\tau_2)) + \\ &+ \sum_{m=1}^{M} (-0.001 \cdot ln(100 w)) \end{aligned} \tag{3}$$

Summarization is carried out for all synapses in the model.

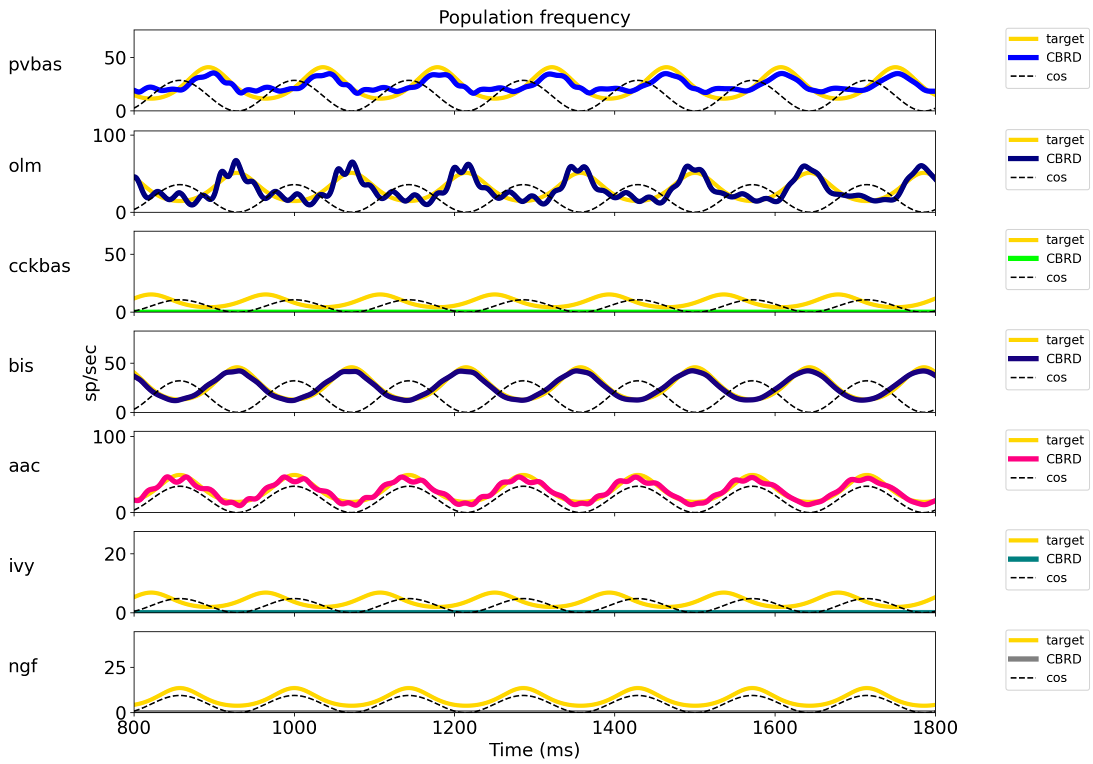


Figure 1: Optimization results of the model with nonplastic synapses. For each population, plots show the target function, the population spike rate obtained with the CBRD approach. One second of simulation is shown after stabilization of the model dynamic mode. The notation of neurons is similar to Fig. 1 of the article.

#### 2 Tables of optimal parameters

Optimal parameters for simulations with plastic synapses.

Table 1: $I_{ext}, \mu A/cm^2$

| | pvbas | olm | cckbas | bis | aac | ivy | ngf |
|---|---|---|---|---|---|---|---|
| Iext | 0.12 | 0.26 | -0.06 | 0.33 | 0.2 | -0.06 | 0.3 |

Table 2: $g_{syn,max}$ $mS$

| | Postsynaptic | | | | | | |
|---|---|---|---|---|---|---|---|
| Presynaptic | pvbas | olm | cckbas | bis | aac | ivy | ngf |
| ca3pyr | 1.14 | 0.68 | 1.14 | 1.14 | 1.16 | 1.16 | 1.63 |
| ca1pyr | 1.47 | 1.82 | 1.83 | 2.22 | 1.61 | 2.08 | - |
| ec3 | 1.38 | - | 1.9 | - | 1.61 | 1.73 | 1.77 |
| pvbas | 3.26 | - | 3.09 | 2.88 | 3.58 | 3.23 | - |
| olm | 1.4 | - | 1.84 | - | 1.69 | - | 1.83 |
| cckbas | 1.13 | - | 1.48 | 1.42 | 1.3 | 1.62 | - |
| bis | 1.39 | 1.16 | 1.69 | 1.61 | 1.43 | 1.64 | - |
| aac | - | - | - | - | - | - | - |
| ivy | 1.48 | 1.33 | 1.6 | 1.46 | 1.65 | 1.62 | 1.61 |
| ngf | 1.42 | - | 1.64 | - | 1.45 | - | 1.64 |

Table 3: w

| | Postsynaptic | | | | | | |
|---|---|---|---|---|---|---|---|
| Presynaptic | pvbas | olm | cckbas | bis | aac | ivy | ngf |
| ca3pyr | 6.57 | 4.26 | 4.41 | 4.91 | 6.38 | 4.6 | 5.02 |
| ca1pyr | 6.3 | 5.83 | 4.65 | 5.15 | 5.63 | 4.33 | - |
| ec3 | 1.63 | - | 1.77 | - | 1.85 | 1.82 | 1.89 |
| pvbas | 0.09 | - | 0.06 | 0.1 | 0.14 | 0.05 | - |
| olm | 0.12 | - | 0.12 | - | 0.11 | - | 0.23 |
| cckbas | 0.06 | - | 0.15 | 0.02 | 0.03 | 0.1 | - |
| bis | 0.04 | 0.15 | 0.16 | 0.2 | 0.11 | 0.15 | - |
| aac | - | - | - | - | - | - | - |
| ivy | 0.08 | 0.03 | 0.09 | 0.03 | 0.18 | 0.16 | 0.15 |
| ngf | 0.1 | - | 0.11 | - | 0.04 | - | 0.17 |

Table 4: $\tau_d, ms$

| | Postsynaptic | | | | | | |
|---|---|---|---|---|---|---|---|
| Presynaptic | pvbas | olm | cckbas | bis | aac | ivy | ngf |
| ca3pyr | 5.61 | 5.34 | 4.78 | 6.14 | 5.61 | 6.51 | 4.18 |
| ca1pyr | 3.25 | 3.21 | 2.74 | 3.45 | 3.4 | 3.82 | - |
| ec3 | 3.89 | - | 3.5 | - | 3.8 | 4.27 | 4.34 |
| pvbas | 3.79 | - | 4.19 | 5.07 | 4.06 | 5.04 | - |
| olm | 5.63 | - | 5.52 | - | 5.89 | - | 6.99 |
| cckbas | 7.01 | - | 6.69 | 8.18 | 7.39 | 8.41 | - |
| bis | 7.46 | 7.93 | 8.03 | 10.21 | 8.01 | 10.68 | - |
| aac | - | - | - | - | - | - | - |
| ivy | 6.57 | 7.34 | 7.01 | 9.03 | 7.05 | 9.37 | 8.86 |
| ngf | 6.5 | - | 6.25 | - | 6.64 | - | 8.84 |

Table 5: $\tau_f, ms$

| | Postsynaptic | | | | | | |
|---|---|---|---|---|---|---|---|
| Presynaptic | pvbas | olm | cckbas | bis | aac | ivy | ngf |
| ca3pyr | 29.78 | 38.38 | 61.44 | 27.59 | 31.54 | 21.98 | 50.32 |
| ca1pyr | 76.9 | 106.95 | 200.09 | 45.85 | 68.72 | 29.95 | - |
| ec3 | 38.31 | - | 97.25 | - | 43.02 | 50.31 | 50.31 |
| pvbas | 15.09 | - | 27.37 | 21.41 | 17.51 | 17.37 | - |
| olm | 16.62 | - | 36.47 | - | 19.45 | - | 20.7 |
| cckbas | 53.25 | - | 89.24 | 49.12 | 68.12 | 39.24 | - |
| bis | 12.43 | 17.63 | 40.15 | 14.36 | 15.79 | 11.64 | - |
| aac | - | - | - | - | - | - | - |
| ivy | 14.36 | 20.69 | 35.03 | 16.88 | 18.31 | 13.43 | 25.56 |
| ngf | 20.48 | - | 40.38 | - | 23.47 | - | 25.58 |

Table 6: $\tau_r, ms$

| | Postsynaptic | | | | | | |
|---|---|---|---|---|---|---|---|
| Presynaptic | pvbas | olm | cckbas | bis | aac | ivy | ngf |
| ca3pyr | 440.07 | 358.57 | 330.99 | 369.46 | 388.4 | 419.17 | 345.25 |
| ca1pyr | 327.26 | 201.9 | 170.37 | 242.43 | 295.47 | 294.63 | - |
| ec3 | 363.56 | - | 274.88 | - | 331.47 | 345.21 | 345.24 |
| pvbas | 635.7 | - | 576.53 | 584.65 | 596.75 | 598.76 | - |
| olm | 650.22 | - | 527.62 | - | 602.4 | - | 578.02 |
| cckbas | 752.48 | - | 638.85 | 628.55 | 663.69 | 653.68 | - |
| bis | 776.48 | 717.22 | 546.21 | 710.0 | 738.36 | 760.08 | - |
| aac | - | - | - | - | - | - | - |
| ivy | 736.06 | 669.69 | 569.35 | 641.27 | 679.4 | 688.89 | 553.77 |
| ngf | 598.63 | - | 500.4 | - | 547.04 | - | 553.66 |

Table 7: $U_{inc}$

| | Postsynaptic | | | | | | |
|---|---|---|---|---|---|---|---|
| Presynaptic | pvbas | olm | cckbas | bis | aac | ivy | ngf |
| ca3pyr | 0.37 | 0.0 | 0.17 | 0.22 | 0.38 | 0.16 | 0.14 |
| ca1pyr | 0.25 | 0.32 | 0.25 | 0.29 | 0.27 | 0.18 | - |
| ec3 | 0.14 | - | 0.28 | - | 0.3 | 0.3 | 0.21 |
| pvbas | 0.29 | - | 0.27 | 0.2 | 0.3 | 0.31 | - |
| olm | 0.29 | - | 0.31 | - | 0.29 | - | 0.36 |
| cckbas | 0.34 | - | 0.26 | 0.23 | 0.34 | 0.2 | - |
| bis | 0.22 | 0.26 | 0.35 | 0.35 | 0.32 | 0.34 | - |
| aac | - | - | - | - | - | - | - |
| ivy | 0.29 | 0.19 | 0.32 | 0.23 | 0.38 | 0.28 | 0.31 |
| ngf | 0.29 | - | 0.27 | - | 0.21 | - | 0.28 |
